# Wider spread of excitatory neuron influence in association cortex than sensory cortex

**DOI:** 10.1101/2024.02.20.581200

**Authors:** Christine F. Khoury, Michael Ferrone, Caroline A. Runyan

**Affiliations:** Department of Neuroscience; Center for the Neural Basis of Cognition, University of Pittsburgh, Pittsburgh, PA

**Keywords:** Inhibitory interneurons, cortical circuits, somatostatin, optogenetics

## Abstract

The basic structure of local cortical circuits, including the composition of cell types, is highly conserved across the cortical processing hierarchy. However, computational roles and the spatial and temporal properties of population activity differ fundamentally in sensory-level and association-level areas. In primary sensory cortex, the timescale of population activity is shorter and pairwise correlations decay more rapidly over distance between neurons, supporting a population code that is suited to encoding rapidly fluctuating sensory stimuli. In association cortex, the timescale of population activity is longer, and pairwise correlations are stronger over wider distances, a code that is suited to holding information in memory and driving behavior. Here, we tested whether these differences in population codes could potentially be explained by intrinsic differences in local network structure. We targeted single excitatory neurons optogenetically, while monitoring the surrounding ongoing population activity in sensory (auditory cortex) and association (posterior parietal cortex) areas in mice. While the temporal impacts of these perturbations were similar across regions, the spatial spread of excitatory influence was wider in association cortex than in sensory cortex. Our findings suggest that differences in recurrent connectivity could contribute to the different properties of population codes in sensory and association cortex, and imply that circuit models of cortical function should be tailored to the properties specific to individual regions.

**Significance statement:** Cell-type-specific functional interactions and connectivity patterns have largely been studied in sensory cortex. Yet the properties of local network activity differ dramatically across the cortical hierarchy, possibly due to differences in intrinsic connectivity patterns. Here, we compared the functional impacts of individual excitatory neurons on local population activity, finding differences in the spatial spread of excitatory influence across regions. Our findings suggest that the structure of local networks differs across the cortical processing hierarchy, and these differences should be considered in circuit models of processes such as decision-making and working memory.

## Introduction

Cortical circuits consist of a rich diversity of cell types, which interact to transform incoming signals by shaping receptive field properties (1), controlling response gain (2–4), and amplifying (5, 6) or sparsening responses to stimuli (7, 8). Circuit dissection experiments in sensory cortex have begun to define the functional and spatial properties of these local interactions and their corresponding computational impacts (7–12).

Excitatory neurons participate in functionally specific recurrent subnetworks in sensory cortex, where neurons with similar response properties tend to form stronger, bidirectional synaptic connections (5, 6, 13). In vivo, the network-level impact of single or small groups of excitatory neurons on sensory responses can lead to either signal amplification or sparsening, possibly depending on stimulus context, network state, and stimulation parameters (7, 8). Therefore, recurrent excitatory interactions seem to have state-dependent functions in sensory cortex.

Inhibitory interneurons are key in modulating local network state, and thus the impacts of feedforward and recurrent excitation. Interneurons can be subdivided into several specific classes that densely interconnect with the local population (9, 12, 14–16), according to different motifs (11, 17–19), some of which include functionally specific connections between excitatory and inhibitory neurons (11, 14). Through these cell-type-specific interactions, inhibitory interneurons can profoundly shift population activity dynamics and also shape the response properties of individual neurons.

For example, somatostatin-expressing interneurons (SOM) inhibit other inhibitory interneurons and also directly inhibit excitatory pyramidal cells (19, 20), refining receptive fields (1, 3, 21), decorrelating population activity at slow timescales (22), and modulating high frequency oscillations (23). SOM neurons tend to be highly coordinated as a population (24–26), especially in association-level cortex, where shared variability has a wider spatial scale than in sensory cortex (24, 27). Potential sources of this coordination among SOM neurons are increased density of excitatory recurrent connections, gap junctions among SOM neurons (18, 28), or modulation of the SOM population by extrinsic inputs.

Cellular and synaptic-level circuit manipulations in the cortex have largely been focused on sensory regions, and so it is not yet known how the spatial and functional structure of local circuit connectivity may differ across cortical regions with different computational goals. For example, in sensory cortex, topographically organized inputs from the thalamus generate the initial tuning underlying sensory representations (29). In association-level areas such as the posterior parietal cortex (PPC), sensory inputs across several modalities are combined (30–32) toward sensorimotor decisions (33–36), perhaps requiring larger scale local circuits.

We hypothesized that differing density and spatial scales of local excitatory connectivity could be a key underlying mechanism explaining differing spatial and temporal scales of correlated variability across cortex (24, 27, 37) and across cell types (24). If this is the case, we would expect the activity of single excitatory neurons to influence a wider local population in association cortex than in sensory cortex. We targeted photostimulation to individual excitatory neurons in auditory cortex (AC) and posterior parietal cortex (PPC) while monitoring activity in neighboring excitatory (E) and SOM neurons. Our results suggest that E cells in PPC do influence the neighboring population of E and SOM neurons over a wider distance than in AC, likely contributing to the differing scales of population activity patterns across cortex.

## Results

### Mapping the influence of excitatory neurons on the local population in vivo

Our goal was to compare the spatial scale of local excitatory network influence in sensory and association cortex. We expressed a red shifted opsin (C1V1) in excitatory (E) neurons, a red fluorophore (tdTomato) in SOM neurons, and a green calcium indicator (GCaMP6f) in all neurons in both auditory cortex (AC) and posterior parietal cortex (PPC) of 10 SOM-Cre x ai14 mice (Figure 1A-B). During imaging sessions (25 AC sessions and 30 PPC sessions), mice ran voluntarily on a spherical treadmill. Neurons expressing tdTomato were considered SOM+, and neurons not expressing tdTomato were considered Non-SOM, the majority of which are excitatory. To map the influence of E neurons on local SOM and Non-SOM neurons in AC and PPC, we could then perform spiral scans of a 1045 nm laser beam over individual E neurons to trigger action potentials (8, 38) while simultaneously raster scanning a 920 nm laser beam to monitor spike-related calcium activity in GCaMP-expressing neurons. In a typical imaging session, we targeted spirals (100 ms in duration) to ∼30 individual E neurons for at least 100 trial repeats, in pseudorandom order, with 1 s between stimulations. Each field of view contained 14.8±6.6 SOM neurons and 222.9±96.0 Non-SOM neurons in AC and PPC, and 22.6±6.9 out of 30.6±7.4 targets were successfully stimulated. All imaging and stimulations were targeted to the same z-depths from the cortical surface (120-300 μm).

**Figure 1:**
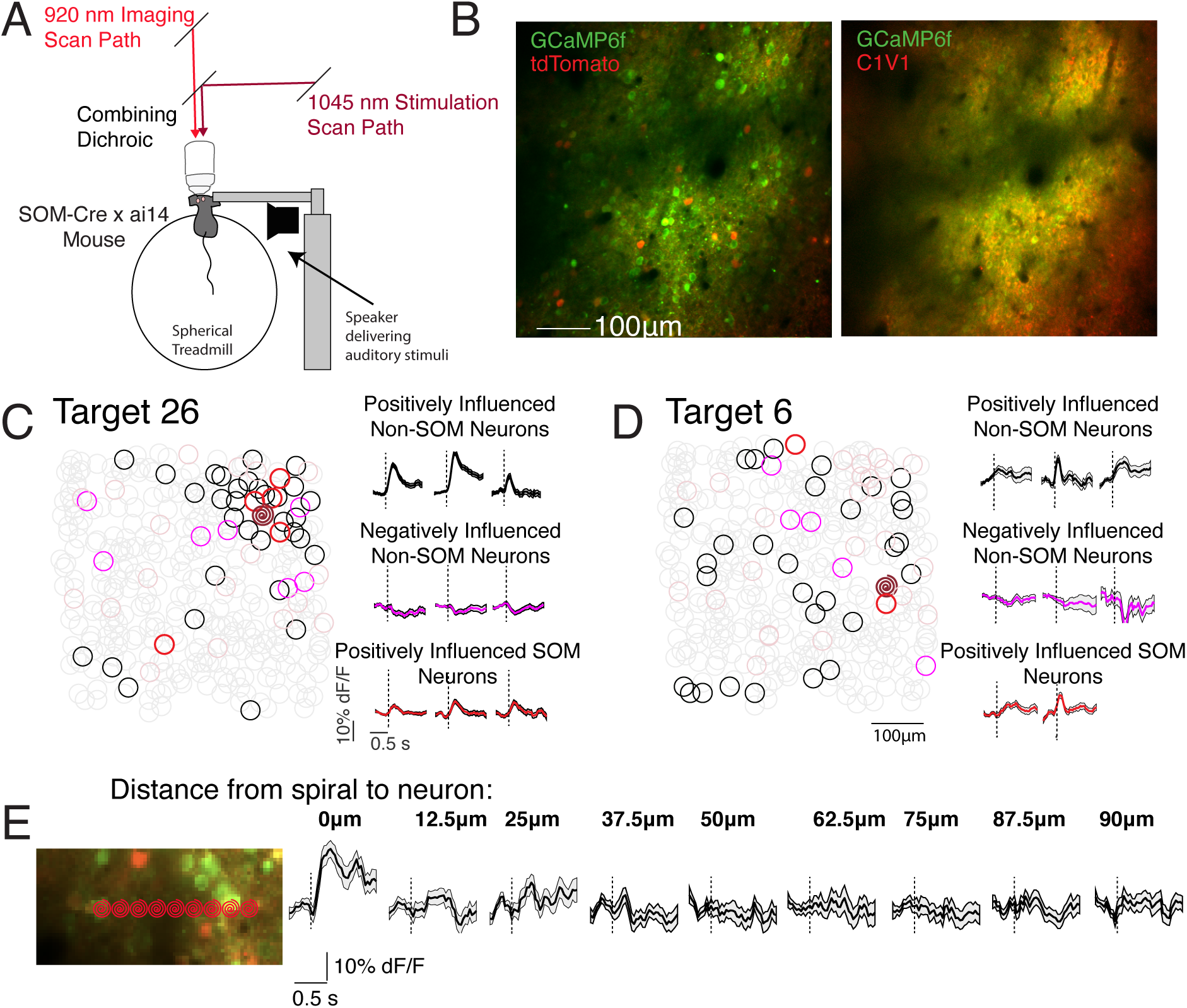
Mapping the influence of E neurons in vivo. **(A)** SOM-Cre x ai14 mice ran voluntarily on a spherical treadmill throughout experiments. A speaker was used to present pure tones to characterize auditory responses. Excitation for imaging was provided by a laser tuned to 920 nm. Excitation for stimulation was provided by a 1045 nm laser that was controlled by an independent scan path. **(B)** An excitatory opsin (C1V1) was virally expressed in excitatory neurons. tdTomato was expressed transgenically in SOM neurons, and GCaMP was expressed in all neurons virally, in SOM-Cre x ai14 mice. Left: Example field of view imaged at 920 nm, visualizing GCaMP (green) and tdTomato (red). Right: Same field of view, imaged at 780 nm, where the mRuby2 (red, also expressed with C1V1) but not tdTomato can be visualized. **(C)** Left: Example influence map obtained from the field of view in B. Excited neuron’s location indicated by the red ‘spiral’. Significantly positively influenced neurons are circled in black, negatively influenced neurons in magenta. Right: Example trial averaged responses of significantly positively influenced E cells, negatively influenced E cells, and positively influenced SOM cells when this neuron was stimulated. **(D)** As in C, for another neuron that was targeted in the same session and field of view. **(E)** Control experiment to determine the spatial resolution of our stimulation. A neuron was stimulated and then spirals were targeted to adjacent locations, as indicated by the spirals. Right: Trial-averaged responses of the neuron under the left-most ‘spiral’.The neuron was only significantly responsive when the spiral was focused directly on it. We only include other neurons that were at least 25μm away from the spiral in our influence measurements. Accoding to our tests, this is a conservative enough approach because we rarely observe direct stimulation of a neuron even 12.5μm away.

The ‘influence map’ for each successfully stimulated neuron was then assessed. We defined ‘influence’ of a target on another neuron as the neuron’s mean response to stimulation (response during 1 s after the target’s stimulation onset minus response in the ∼300 ms before stimulation onset) across all trials, divided by its standard deviation. On average, a sparse subset of neighboring neurons was positively influenced (7.9% ± 8.7%, defined as influence greater than 99% of trial shuffled data, see Methods, considering only neurons at least 25 μm away from the target), and a smaller subset of local neurons were negatively influenced (3.4%, ±3.5%, response was less than 99% of trial shuffled data) (Figure 1 C-D).

To estimate the spatial specificity of the spiral stimulation, in most sessions we selected a single C1V1-expressing neuron that was on the edge of the viral injection site. We then systematically stimulated the location of the neuron’s soma, and neighboring distances at 12.5 μm increments (Figure 1E). For most targets, significant responses were only evoked when the spirals were targeted within 25 μm of the soma. In subsequent analyses, neurons within 25 μm of the stimulation target were excluded.

### E influence spreads farther in PPC than AC

We successfully stimulated 506 E neurons in AC, testing 2,786,036 potential influences on Non-SOM and 150,282 potential influences on SOM neurons. In PPC we successfully stimulated 577 E neurons, testing 3,111,761 potential influences on Non-SOM neurons and 227,338 potential influences on SOM neurons in PPC. In analysis, to examine the spatial spread of E influence in each area, we centered each targeted neuron, and overlaid and summed the relative spatial positions of significantly positively influenced Non-SOM neurons, normalizing by the total number of targeted neurons. The spatial zone of positively influenced neurons appeared to be more restricted in AC than in PPC (Figure 2A). A similar trend was evident when we mapped the positive influence on SOM neurons (Figure 2B), even with the overall sparser distribution of SOM neurons. Similarly, we mapped negative influence on the local population, to illustrate the spatial spread of targets’ negative impacts on activity. Interestingly, negative influence was more spatially scattered than positive influence, and appeared to spread wider in AC than in PPC (Figure 2C).

**Figure 2:**
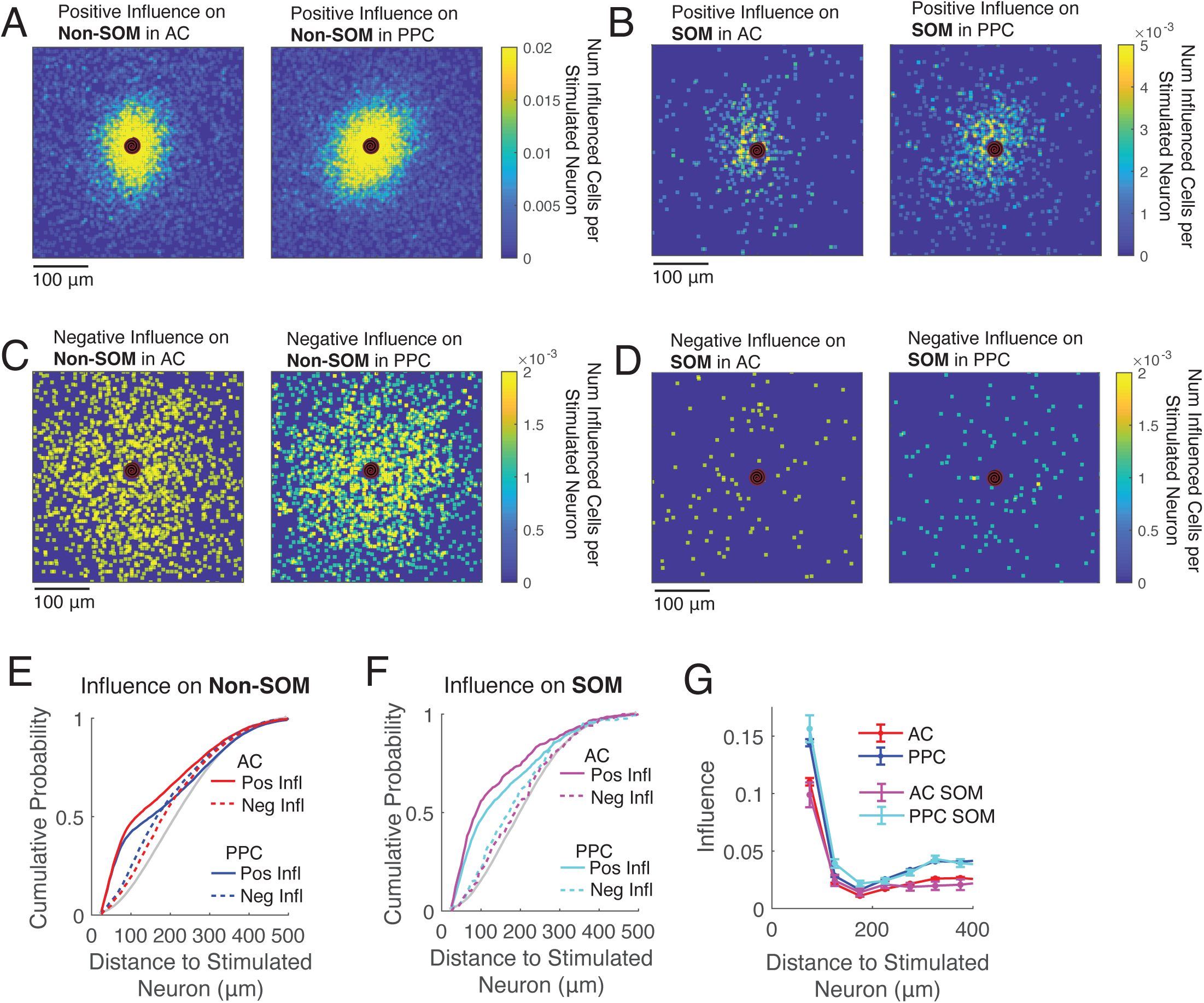
E influence decays with distance in AC and PPC. **(A)** Influence maps on Non-SOM neurons, across all targeted E neurons and all fields of view in AC (left) and PPC (right). Each targeted was centered, and locations of significantly positively influenced neurons set to 1. Then, all 506 AC maps and 577 PPC maps were summed and divided by the total number of maps, so that the color at each pixel location indicates the number of significantly positively influenced Non-SOM neurons per target. **(B)** As in (A) but for positive influence on SOM neurons (n’s are the same as in A). **(C-D)** As in (A-B) but for negative influence. **(E)** Cumulative distributions of the intersomatic distances between the targets and significantly influenced Non-SOM cells, in AC (red) and PPC (blue). Grey: distributions of distances of all Non-SOM neurons to the target. Solid: positive influence, dashed: negative influence. n = 9690, positively influenced AC Non-SOM neurons, 2321 negatively influenced AC Non-SOM neurons, 17120 positively influenced PPC Non-SOM neurons, and 2776 negatively influenced PPC Non-SOM neurons. Total number of AC Non-SOM neurons: 125496. Total number of PPC Non-SOM neurons: 171692. **(F)** As in E, but showing the distributions of distances of SOM neurons to the targets. n = 532 positively influenced AC SOM neurons, 137 negatively influenced AC SOM neurons, 6846 total AC neurons, 1245 positively influenced PPC SOM neurons, 170 negatively influenced PPC SOM neurons, and 12152 total PPC SOM neurons. In all panels, only neurons at least 25 μm from the target were considered. **(G)** Same data as in E-F, but plotting the mean influence on all neurons, binned by distance to the target. See Tables 1-2 for full values and statistics.

**Table 1:** Related to Figure 2.

|  | **<u>Table 1: Related to Figure 2</u>** |  |  |  |
| --- | --- | --- | --- | --- |
|  |  | **Mean Distance to Target** | **Standard Deviation** | **n** |
| **1** | AC All Non-SOM | 226.3 | 110.9 | 125496 |
| **2** | AC Positively Influenced Non-SOM | 162.7 | 123.4 | 9690 |
| **3** | AC Negatively Influenced Non-SOM | 202.7 | 109 | 2321 |
| **4** | AC Un-Influenced Non-SOM | 232.3 | 108 | 113485 |
| **5** | PPC All Non-SOM | 209 | 103.9 | 171692 |
| **6** | PPC Positively Influenced Non-SOM | 159.8 | 120.1 | 17120 |
| **7** | PPC Negatively Influenced Non-SOM | 170.4 | 95.5 | 2776 |
| **8** | PPC Un-Influenced Non-SOM | 215.2 | 100.4 | 151796 |
| **9** | AC All SOM | 210 | 109.9 | 6846 |
| **10** | AC Positively Influenced SOM | 136.3 | 109.6 | 532 |
| **11** | AC Negatively Influenced SOM | 197.9 | 101.8 | 137 |
| **12** | AC Un-Influenced SOM | 219.3 | 106 | 6094 |
| **13** | PPC All SOM | 209 | 105.4 | 12152 |
| **14** | PPC Positively Influenced SOM | 154.8 | 113.9 | 1245 |
| **15** | PPC Negatively Influenced SOM | 187.5 | 103.6 | 170 |
| **16** | PPC Un-Influenced SOM | 217.7 | 101 | 10623 |
| **Comparison\*** | **p** | **Difference** | **effect size** |  |
| 9,13 | 0.3526 | 1.5 | 0.0142 |  |
| 10,14 | 9.99E-04 | -18.6 | -0.1649 |  |
| 11,15 | 0.3644 | 10.4 | 0.1016 |  |
| 1,5 | 9.99E-04 | -17.4 | -0.1625 |  |
| 2,6 | 0.0689 | 2.9 | 0.0237 |  |
| 3,7 | 9.99E-04 | 32.2 | 0.316 |  |
| 1,2 | 9.99E-04 | 63.7 | 0.5695 |  |
| 1,3 | 9.99E-04 | 23.7 | 0.2137 |  |
| 1,4 | 9.99E-04 | -5.9 | -0.0541 |  |
| 2,3 | 9.99E-04 | -40 | -0.3311 |  |
| 3,4 | 9.99E-04 | -29.6 | -0.2742 |  |
| 5,6 | 9.99E-04 | 49.2 | 0.4663 |  |
| 5,7 | 9.99E-04 | 38.5 | 0.3713 |  |
| 5,8 | 9.99E-04 | -6.2 | -0.0611 |  |
| 6,7 | 9.99E-04 | -10.7 | -0.0911 |  |
| 7,8 | 9.99E-04 | -44.8 | -0.4465 |  |
| 1,9 | 9.99E-04 | 14 | 0.1269 |  |
| 2,10 | 9.99E-04 | 26.4 | 0.2152 |  |
| 3,11 | 0.6154 | 4.76 | 0.0438 |  |
| 9,10 | 9.99E-04 | 76 | 0.7003 |  |
| 9,11 | 0.1229 | 14.4 | 0.1328 |  |
| 9,12 | 2.00E-04 | -7 | -6.49E-02 |  |
| 10,11 | 1.00E-04 | -61.6 | -5.71E-01 |  |
| 11,12 | 0.0224 | -21.4 | -2.02E-01 |  |
| 5,13 | 0.0699 | -1.8 | -1.75E-02 |  |
| 6,14 | 0.1518 | 4.9 | 0.0414 |  |
| 7,15 | 0.036 | -17 | -0.1771 |  |
| 13,14 | 1.00E-04 | 56 | 0.532 |  |
| 13,15 | 0.0042 | 23.3 | 0.224 |  |
| 13,16 | 1.00E-04 | -6.9 | -0.0675 |  |
| 14,15 | 9.00E-04 | -32.6 | -0.2893 |  |
| 15,16 | 3.00E-04 | -30.3 | -0.2997 |  |
| *Permutation Test |  |  |  |  |

**Table 2:** ANOVA and Posthoc Tests, Related to Figure 2.

| Group Number | Group Name | Mean Influence | Standard Deviation |
| --- | --- | --- | --- |
| 1 | AC,Non-SOM,60um | 0.257 | 0.002 |
| 2 | PPC,Non-SOM,60um | 0.343 | 0.002 |
| 3 | AC,SOM,60um | 0.233 | 0.008 |
| 4 | PPC,SOM,60um | 0.335 | 0.007 |
| 5 | AC,Non-SOM,90um | 0.095 | 0.002 |
| 6 | PPC,Non-SOM,90um | 0.126 | 0.001 |
| 7 | AC,SOM,90um | 0.090 | 0.007 |
| 8 | PPC,SOM,90um | 0.142 | 0.005 |
| 9 | AC,Non-SOM,120um | 0.029 | 0.002 |
| 10 | PPC,Non-SOM,120um | 0.038 | 0.001 |
| 11 | AC,SOM,120um | 0.036 | 0.006 |
| 12 | PPC,SOM,120um | 0.050 | 0.005 |
| 13 | AC,Non-SOM,150um | 0.012 | 0.001 |
| 14 | PPC,Non-SOM,150um | 0.015 | 0.001 |
| 15 | AC,SOM,150um | 0.010 | 0.006 |
| 16 | PPC,SOM,150um | 0.028 | 0.004 |
| 17 | AC,Non-SOM,180um | 0.012 | 0.001 |
| 18 | PPC,Non-SOM,180um | 0.019 | 0.001 |
| 19 | AC,SOM,180um | 0.019 | 0.006 |
| 20 | PPC,SOM,180um | 0.027 | 0.004 |
| 21 | AC,Non-SOM,210um | 0.016 | 0.001 |
| 22 | PPC,Non-SOM,210um | 0.025 | 0.001 |
| 23 | AC,SOM,210um | 0.012 | 0.006 |
| 24 | PPC,SOM,210um | 0.025 | 0.004 |
| 25 | AC,Non-SOM,240um | 0.020 | 0.001 |
| 26 | PPC,Non-SOM,240um | 0.029 | 0.001 |
| 27 | AC,SOM,240um | 0.027 | 0.006 |
| 28 | PPC,SOM,240um | 0.036 | 0.004 |
| 29 | AC,Non-SOM,270um | 0.021 | 0.001 |
| 30 | PPC,Non-SOM,270um | 0.035 | 0.001 |
| 31 | AC,SOM,270um | 0.017 | 0.006 |
| 32 | PPC,SOM,270um | 0.034 | 0.005 |
| 33 | AC,Non-SOM,300um | 0.025 | 0.001 |
| 34 | PPC,Non-SOM,300um | 0.037 | 0.001 |
| 35 | AC,SOM,300um | 0.016 | 0.007 |
| 36 | PPC,SOM,300um | 0.037 | 0.005 |
| 37 | AC,Non-SOM,330um | 0.026 | 0.002 |
| 38 | PPC,Non-SOM,330um | 0.043 | 0.001 |
| 39 | AC,SOM,330um | 0.019 | 0.007 |
| 40 | PPC,SOM,330um | 0.044 | 0.006 |
| 41 | AC,Non-SOM,360um | 0.026 | 0.002 |
| 42 | PPC,Non-SOM,360um | 0.042 | 0.002 |
| 43 | AC,SOM,360um | 0.018 | 0.008 |
| 44 | PPC,SOM,360um | 0.038 | 0.006 |
| 45 | AC,Non-SOM,390um | 0.028 | 0.002 |
| 46 | PPC,Non-SOM,390um | 0.046 | 0.002 |
| 47 | AC,SOM,390um | 0.023 | 0.009 |
| 48 | PPC,SOM,390um | 0.039 | 0.008 |
| 49 | AC,Non-SOM,420um | 0.028 | 0.002 |
| 50 | PPC,Non-SOM,420um | 0.042 | 0.002 |
| 51 | AC,SOM,420um | 0.025 | 0.011 |
| 52 | PPC,SOM,420um | 0.040 | 0.009 |
| 53 | AC,Non-SOM,450um | 0.024 | 0.003 |
| 54 | PPC,Non-SOM,450um | 0.045 | 0.003 |
| 55 | AC,SOM,450um | 0.033 | 0.014 |
| 56 | PPC,SOM,450um | 0.044 | 0.011 |
| 57 | AC,Non-SOM,480um | 0.029 | 0.003 |
| 58 | PPC,Non-SOM,480um | 0.047 | 0.004 |
| 59 | AC,SOM,480um | 0.020 | 0.018 |
| 60 | PPC,SOM,480um | 0.042 | 0.014 |
| 61 | AC,Non-SOM,510um | 0.027 | 0.003 |
| 62 | PPC,Non-SOM,510um | 0.062 | 0.004 |
| 63 | AC,SOM,510um | 0.028 | 0.017 |
| 64 | PPC,SOM,510um | 0.030 | 0.013 |

Posthoc Multiple Comparisons Results (Less relevant comparisons removed for brevity)

| Group A | Group B | Lower Bound | Estimated Difference | Upper Bound | p-value |
| --- | --- | --- | --- | --- | --- |
| 1 | 2 | -0.098 | -0.087 | -0.075 | 9.59E-06 |
| 1 | 3 | -0.013 | 0.023 | 0.059 | 0.9453525921 |
| 1 | 4 | -0.107 | -0.078 | -0.050 | 9.59E-06 |
| 1 | 5 | 0.150 | 0.162 | 0.173 | 9.59E-06 |
| 1 | 6 | 0.120 | 0.130 | 0.141 | 9.59E-06 |
| 1 | 7 | 0.138 | 0.167 | 0.196 | 9.59E-06 |
| 1 | 8 | 0.091 | 0.114 | 0.138 | 9.59E-06 |
| 2 | 3 | 0.074 | 0.110 | 0.145 | 9.59E-06 |
| 2 | 4 | -0.020 | 0.008 | 0.036 | 1 |
| 2 | 5 | 0.238 | 0.248 | 0.258 | 9.59E-06 |
| 2 | 6 | 0.208 | 0.217 | 0.226 | 9.59E-06 |
| 2 | 7 | 0.225 | 0.253 | 0.282 | 9.59E-06 |
| 2 | 8 | 0.178 | 0.201 | 0.224 | 9.59E-06 |
| 2 | 9 | 0.305 | 0.315 | 0.324 | 9.59E-06 |
| 3 | 4 | -0.145 | -0.102 | -0.058 | 9.59E-06 |
| 3 | 5 | 0.103 | 0.139 | 0.174 | 9.59E-06 |
| 3 | 6 | 0.072 | 0.107 | 0.142 | 9.59E-06 |
| 3 | 7 | 0.099 | 0.144 | 0.188 | 9.59E-06 |
| 3 | 8 | 0.050 | 0.091 | 0.132 | 9.59E-06 |
| 3 | 9 | 0.170 | 0.205 | 0.240 | 9.59E-06 |
| 4 | 5 | 0.212 | 0.240 | 0.268 | 9.59E-06 |
| 4 | 6 | 0.181 | 0.209 | 0.236 | 9.59E-06 |
| 4 | 7 | 0.207 | 0.245 | 0.284 | 9.59E-06 |
| 4 | 8 | 0.158 | 0.193 | 0.228 | 9.59E-06 |
| 5 | 6 | -0.040 | -0.031 | -0.022 | 9.59E-06 |
| 5 | 7 | -0.023 | 0.005 | 0.034 | 1 |
| 5 | 8 | -0.070 | -0.047 | -0.024 | 9.59E-06 |
| 6 | 7 | 0.008 | 0.036 | 0.065 | 0.0002 |
| 6 | 8 | -0.038 | -0.016 | 0.007 | 0.8320 |
| 6 | 9 | 0.089 | 0.098 | 0.106 | 9.59E-06 |
| 7 | 8 | -0.088 | -0.052 | -0.017 | 1.15E-05 |
| 9 | 10 | -0.017 | -0.009 | -0.001 | 0.004083 |
| 9 | 11 | -0.034 | -0.008 | 0.019 | 1.000000 |
| 9 | 12 | -0.041 | -0.022 | -0.002 | 0.011075 |
| 9 | 13 | 0.008 | 0.017 | 0.026 | 0.000010 |
| 9 | 14 | 0.005 | 0.013 | 0.021 | 0.000010 |
| 9 | 15 | -0.007 | 0.018 | 0.043 | 0.784965 |
| 9 | 16 | -0.019 | 0.001 | 0.020 | 1.000000 |
| 10 | 11 | -0.025 | 0.002 | 0.028 | 1.000000 |
| 10 | 12 | -0.032 | -0.012 | 0.007 | 0.960780 |
| 10 | 13 | 0.019 | 0.026 | 0.034 | 0.000010 |
| 10 | 14 | 0.016 | 0.023 | 0.030 | 0.000010 |
| 10 | 15 | 0.003 | 0.027 | 0.052 | 0.008952 |
| 10 | 16 | -0.009 | 0.010 | 0.029 | 0.999184 |
| 11 | 12 | -0.046 | -0.014 | 0.018 | 0.999997 |
| 11 | 13 | -0.002 | 0.025 | 0.051 | 0.133744 |
| 11 | 14 | -0.005 | 0.021 | 0.047 | 0.496362 |
| 11 | 15 | -0.010 | 0.026 | 0.061 | 0.769850 |
| 11 | 16 | -0.023 | 0.008 | 0.040 | 1.000000 |
| 12 | 13 | 0.019 | 0.039 | 0.058 | 9.59E-06 |
| 12 | 14 | 0.016 | 0.035 | 0.054 | 9.59E-06 |
| 12 | 15 | 0.009 | 0.040 | 0.070 | 0.000207 |
| 12 | 16 | -0.004 | 0.022 | 0.049 | 0.318831 |
| 12 | 64 | -0.035 | 0.021 | 0.076 | 1.000000 |
| 13 | 14 | -0.011 | -0.004 | 0.004 | 0.999965 |
| 13 | 15 | -0.024 | 0.001 | 0.026 | 1.000000 |
| 13 | 16 | -0.036 | -0.016 | 0.003 | 0.324978 |
| 14 | 15 | -0.020 | 0.005 | 0.029 | 1.000000 |
| 14 | 16 | -0.032 | -0.013 | 0.006 | 0.900593 |
| 15 | 16 | -0.048 | -0.017 | 0.013 | 0.99428 |
| 15 | 17 | -0.026 | -0.001 | 0.024 | 1.00000 |
| 17 | 18 | -0.014 | -0.007 | 0.000 | 0.12237 |
| 17 | 19 | -0.032 | -0.008 | 0.017 | 1.00000 |
| 17 | 20 | -0.033 | -0.015 | 0.004 | 0.47611 |
| 18 | 19 | -0.025 | -0.001 | 0.023 | 1.00000 |
| 18 | 20 | -0.026 | -0.008 | 0.010 | 1.00000 |
| 18 | 21 | -0.005 | 0.002 | 0.010 | 1.00000 |
| 19 | 20 | -0.037 | -0.007 | 0.022 | 1.00000 |
| 21 | 22 | -0.016 | -0.009 | -0.001 | 0.00198 |
| 21 | 23 | -0.020 | 0.005 | 0.029 | 1.00000 |
| 21 | 24 | -0.027 | -0.008 | 0.010 | 0.99999 |
| 22 | 23 | -0.011 | 0.013 | 0.037 | 0.99779 |
| 22 | 24 | -0.018 | 0.000 | 0.019 | 1.00000 |
| 23 | 24 | -0.043 | -0.013 | 0.017 | 1.00000 |
| 25 | 26 | -0.016 | -0.009 | -0.001 | 0.00233 |
| 25 | 27 | -0.033 | -0.007 | 0.019 | 1.00000 |
| 25 | 28 | -0.035 | -0.016 | 0.003 | 0.31873 |
| 26 | 27 | -0.024 | 0.002 | 0.028 | 1.00000 |
| 26 | 28 | -0.026 | -0.007 | 0.011 | 1.00000 |
| 27 | 28 | -0.040 | -0.009 | 0.022 | 1.00000 |
| 29 | 30 | -0.021 | -0.014 | -0.006 | 0.00001 |
| 29 | 31 | -0.022 | 0.004 | 0.030 | 1.00000 |
| 29 | 32 | -0.032 | -0.013 | 0.007 | 0.93777 |
| 30 | 31 | -0.008 | 0.018 | 0.044 | 0.87804 |
| 30 | 32 | -0.018 | 0.001 | 0.020 | 1.00000 |
| 31 | 32 | -0.048 | -0.017 | 0.015 | 0.99907 |
| 33 | 34 | -0.021 | -0.013 | -0.004 | 0.00001 |
| 33 | 35 | -0.020 | 0.009 | 0.037 | 1.00000 |
| 33 | 36 | -0.032 | -0.012 | 0.008 | 0.99095 |
| 34 | 35 | -0.007 | 0.021 | 0.050 | 0.69408 |
| 34 | 36 | -0.020 | 0.001 | 0.021 | 1.00000 |
| 35 | 36 | -0.055 | -0.021 | 0.013 | 0.98091 |
| 37 | 38 | -0.026 | -0.017 | -0.008 | 0.00001 |
| 37 | 39 | -0.024 | 0.006 | 0.037 | 1.00000 |
| 37 | 40 | -0.042 | -0.018 | 0.005 | 0.56449 |
| 38 | 39 | -0.007 | 0.023 | 0.054 | 0.60811 |
| 38 | 40 | -0.025 | -0.002 | 0.022 | 1.00000 |
| 39 | 40 | -0.062 | -0.025 | 0.012 | 0.91682 |
| 41 | 42 | -0.026 | -0.016 | -0.006 | 0.00001 |
| 41 | 43 | -0.027 | 0.008 | 0.042 | 1.00000 |
| 41 | 44 | -0.038 | -0.012 | 0.014 | 0.99999 |
| 42 | 43 | -0.011 | 0.024 | 0.058 | 0.87381 |
| 42 | 44 | -0.022 | 0.004 | 0.030 | 1.00000 |
| 42 | 45 | 0.004 | 0.014 | 0.025 | 0.00007 |
| 42 | 46 | -0.014 | -0.004 | 0.007 | 1.00000 |
| 43 | 44 | -0.062 | -0.020 | 0.023 | 0.99998 |
| 43 | 45 | -0.044 | -0.010 | 0.025 | 1.00000 |
| 43 | 46 | -0.062 | -0.028 | 0.007 | 0.52356 |
| 44 | 45 | -0.016 | 0.010 | 0.036 | 1.00000 |
| 44 | 46 | -0.034 | -0.008 | 0.018 | 1.00000 |
| 45 | 46 | -0.029 | -0.018 | -0.007 | 0.00001 |

To quantify potential differences in the spatial spread of positive and negative influence by E cells in AC and PPC, we measured the inter-somatic distance from the target to each neuron in the field of view. We then compared the distributions of inter-somatic distances of significantly positively and negatively influenced Non-SOM neurons. In both regions, positively influenced neurons tended to be closer to the targeted neurons than both negatively influenced and the full distribution of imaged neurons (p = 0.00099; Figure 2E; see Table 1 for full values and statistics). Similar trends appeared in the SOM population (Figure 2F; Table 1).

To test for effects of brain region, cell type, and inter-somatic distance on the targets’ influence, we performed a three-way ANOVA (Figure 2G; results reported in Table 2). There were significant main effects of brain area and inter-somatic distance (p<0.0001), and significant interactions between brain area and inter-somatic distance (p<0.0001) and between cell type and distance (p = 0.0072). Posthoc multiple comparisons tests revealed greater influence on PPC SOM than AC SOM neurons at distances less than 100 μm (p = 9.59E-06, Tukey-Kramer test for multiple comparisons) and greater influence on PPC than AC Non-SOM neurons at distances less than 100 μm (p = 9.59E-06, see Table 2 for full list of statistical comparisons – Tukey-Kramer test for multiple comparisons).

### The effects of E stimulation were overall sparse, but more positive in the PPC population than AC

We hypothesized that another contributor to the greater level of shared variability in the PPC population could be a higher density of recurrent excitatory synapses, compared to AC. While our method does not have the temporal resolution to identify monosynaptic relationships, we reasoned that an overall higher density of recurrent connections may lead to a higher density of influenced neurons. For each stimulated neuron, we identified a ‘stimulus axis’ in high dimensional space, which is the axis in this space that connects the population’s average prestimulus activity to the population’s response to the stimulation. When we projected population activity onto this axis, a robust response was revealed (Figure 3A). Next, we examined the distribution of weights of the non-stimulated neurons in the stimulus axes. The mean stimulus axis weight was higher and more positive in PPC than AC (AC: 0.018±0.080, PPC: 0.026±0.089, p = 1.00E-04, permutation test), suggesting that while the density of E influence in the two regions is very sparse, more neurons are positively influenced in PPC than in AC (Figure 3B-D).

**Figure 3:**
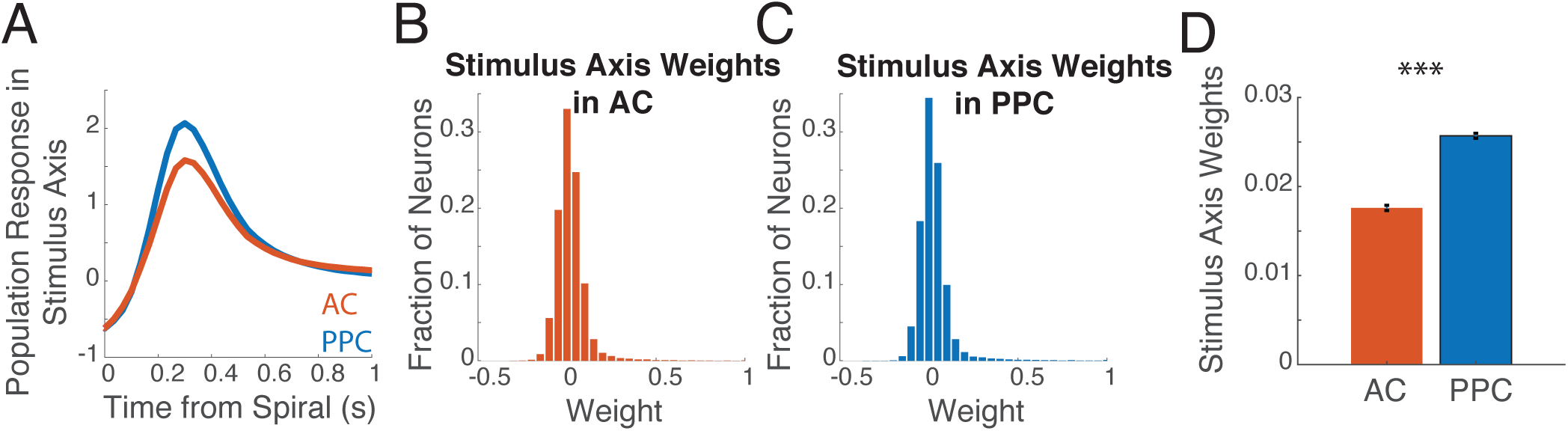
E influence on the local population is generally sparse, but is more positive in PPC than in AC. **(A)** A ‘stimulus axis’ was defined in high dimensional space, that best aligned with the population’s response to the target stimulation. The mean response in the stimulus axis (defined independently for each target stimulus), is plotted for AC and PPC. n = 506 AC populations and 577 PPC populations. **(B)** The distribution of weights along the stimulus axis, of all AC neurons at least 25 μm from the targets (n = 125496 neurons). **(C)** As in B, for PPC neurons (n = 171692 neurons). **(D)** Same data as in B-C, but plotted as bar plots of the mean stimulus axis weight across all AC and all PPC neurons. Error bars indicate standard error of the mean.. PPC weights were significantly more positive than AC weights (p = 1.00E-04, permutation test).

### Relating influence and pairwise noise correlations

In visual cortex, closely neighboring neurons with positive noise correlations (greater shared variability) have a higher monosynaptic connection probability (6), and greater positive influence in similar conditions as our experiments (8). To test if this is a general feature of local circuits across the cortex, we computed Pearson’s correlation coefficients between the activity traces of every target-neuron pair, using only imaging frames outside of the target neuron’s stimulation trials, and related this r value to the influence of the target on the other neuron. In both AC and PPC, influence on Non-SOM neurons was positively related to pairwise noise correlation for cell pairs with intersomatic distances within 125 μm (Pearson r between influence and pairwise noise correlation between neuron and target: A1 Non-SOM: r = 0.17, p < 0.000001; PPC Non-SOM: r = 0.15, p < 0.000001) (Figure 4A-B). The relationship was flatter for pairs with intersomatic distances greater than 125 μm, which is consistent with results in mouse V1 (8) (A1 Non-SOM: r = 0.01, p = 0.0094; PPC Non-SOM: r = 0.01, p = 0.0021). In contrast, the relationship between pairwise noise correlation and influence on SOM neurons was flat among neurons near the target in AC and PPC (Figure 4C) (AC SOM within 125 μm of the target: r = 0.02, p = 0.53; PPC SOM: r = 0.03, p = 0.18). At farther distances, noise correlations and influence on SOM neurons did have a positive relationship in AC and PPC (AC greater than 125 μm from the target: r = 0.08, p = 0.00014; PPC: r = 0.05, p = 0.00046, Figure 4C-D, Table 3 for full values).

**Figure 4:**
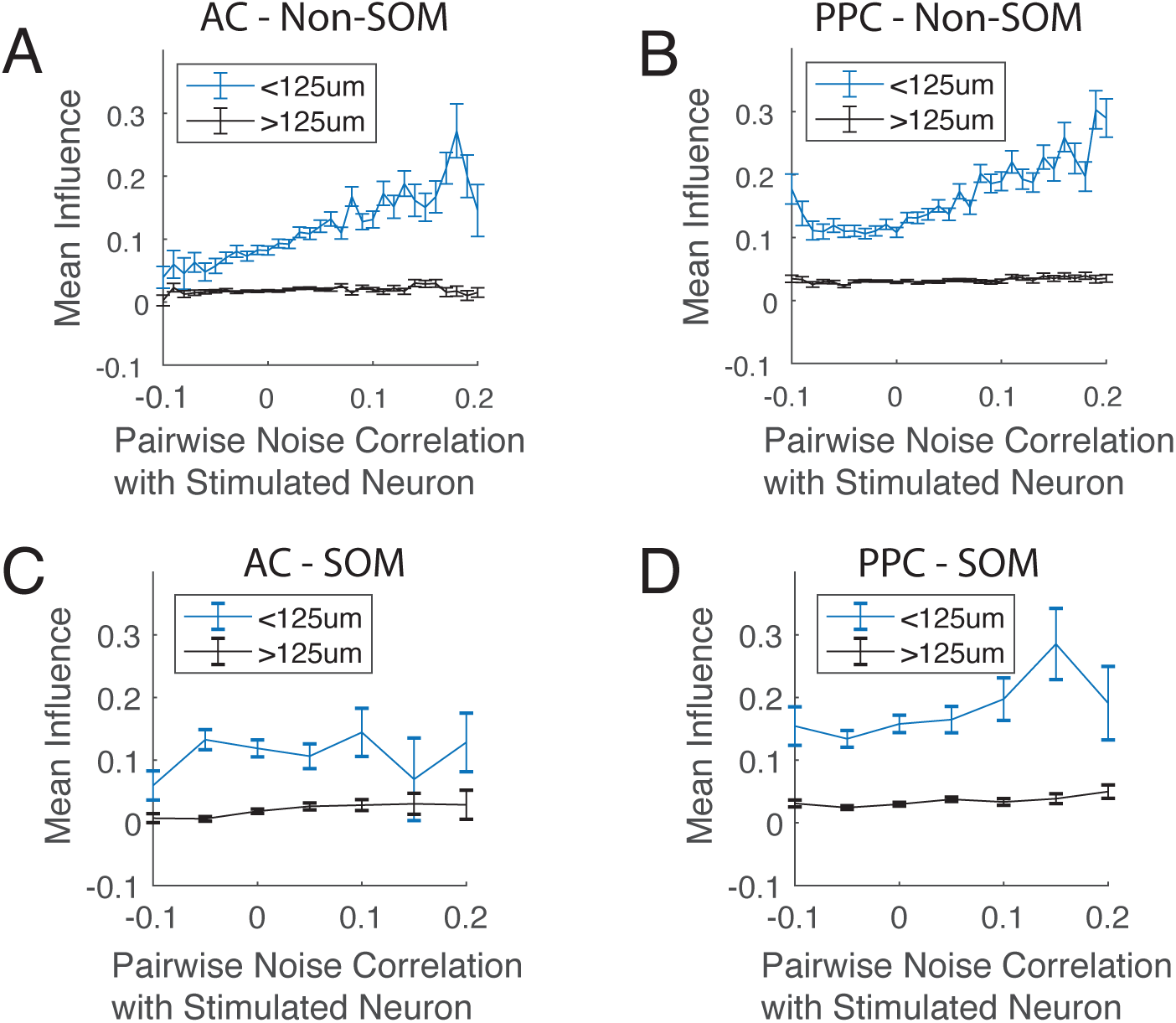
The relationship between E influence and pairwise noise correlation. **(A)** Pairwise Pearson correlations were computed between each target-neuron pair, and related to the target’s influence on the other neuron. Noise correlation and influence were more positively correlated for Non-SOM neurons within 125 μm of the target (blue), than for neurons farther away (black). **(B)** As in A, for PPC Non-SOM neurons. As in AC, neighboring Non-SOM with stronger noise correlations tended to be more positively influenced. **(C)** As in A, for AC SOM neurons. Noise correlations and influence were not positively related in neighboring SOM neurons, but were weakly correlated in SOM neurons at greater distances. **(D)** As in A-C, for PPC SOM neurons. See Table 3 for full values and statistics.

**Table 3:** Related to Figure 4.

| Pearson's r between noise correlation and influence | AC neurons < 125um to target |  | AC neurons > 125 um to target |  | PPC neurons <125um to target |  | PPC neurons >125um to target |  |
| --- | --- | --- | --- | --- | --- | --- | --- | --- |
|  | Non-SOM | SOM | Non-SOM | SOM | Non-SOM | SOM | Non-SOM | SOM |
| **r** | 0.1715 | 0.0215 | 0.0117 | 0.0761 | 0.1489 | 0.0339 | 0.0122 | 0.0511 |
| **p** | 8.17E-96 | 0.5289 | 0.0094 | 1.42E-04 | 5.43E-112 | 0.1854 | 0.0021 | 4.66E-04 |
| **Number of Pairs** | 14446 | 858 | 48875 | 2493 | 22545 | 1530 | 62910 | 4683 |

### E influence on Non-SOM neurons is greater for neurons with similar sound frequency tuning in AC

In V1, E neurons with similar orientation preferences and high signal correlations have a higher monosynaptic connection probability (6), which allows recurrent E connections to provide signal amplification. *In vivo*, single neuron stimulation in V1 can lead to polysynaptic inhibitory influence on neurons with similar tuning (8), a potential mechanism for feature competition. However, stimulation of ensembles of similarly tuned neurons leads to more excitatory effects in similarly tuned neurons (7). To determine whether similar patterns exist in mouse auditory cortex, we played pure tone stimuli, allowing us to characterize the frequency tuning preference for each sound-responsive neuron in AC. We could then relate neurons’ pure tone frequency preferences to their influence.

For this analysis, we considered only AC neurons that were sound responsive and had a preferred sound frequency (71450 target-neuron pairs, Figure 5A). We estimated each neuron’s preferred sound frequency by fitting a Gaussian to the neuron’s trial-averaged responses, and then measured the difference in sound frequency preference between each stimulated target and other neuron in the field of view, in octaves (Figure 5A). Next, we related the influence to binned BF difference, and compared this to a shuffled distribution, where we randomly shuffled the frequency preferences across all neurons in the field of view. In Non-SOM neurons, influence was stronger on neurons with more similar frequency preferences, and weaker for neurons with more different frequency preferences (Pearson’s correlation between influence and relative best frequency was significantly more negative than 500 shuffles, p = 0.014, r = -0.0257, Figure 5B). Influence on SOM neurons, however, was similar to the shuffled distribution, and did not relate to relative tuning (p = 0.12, r = 0.05, Figure 5C).

**Figure 5:**
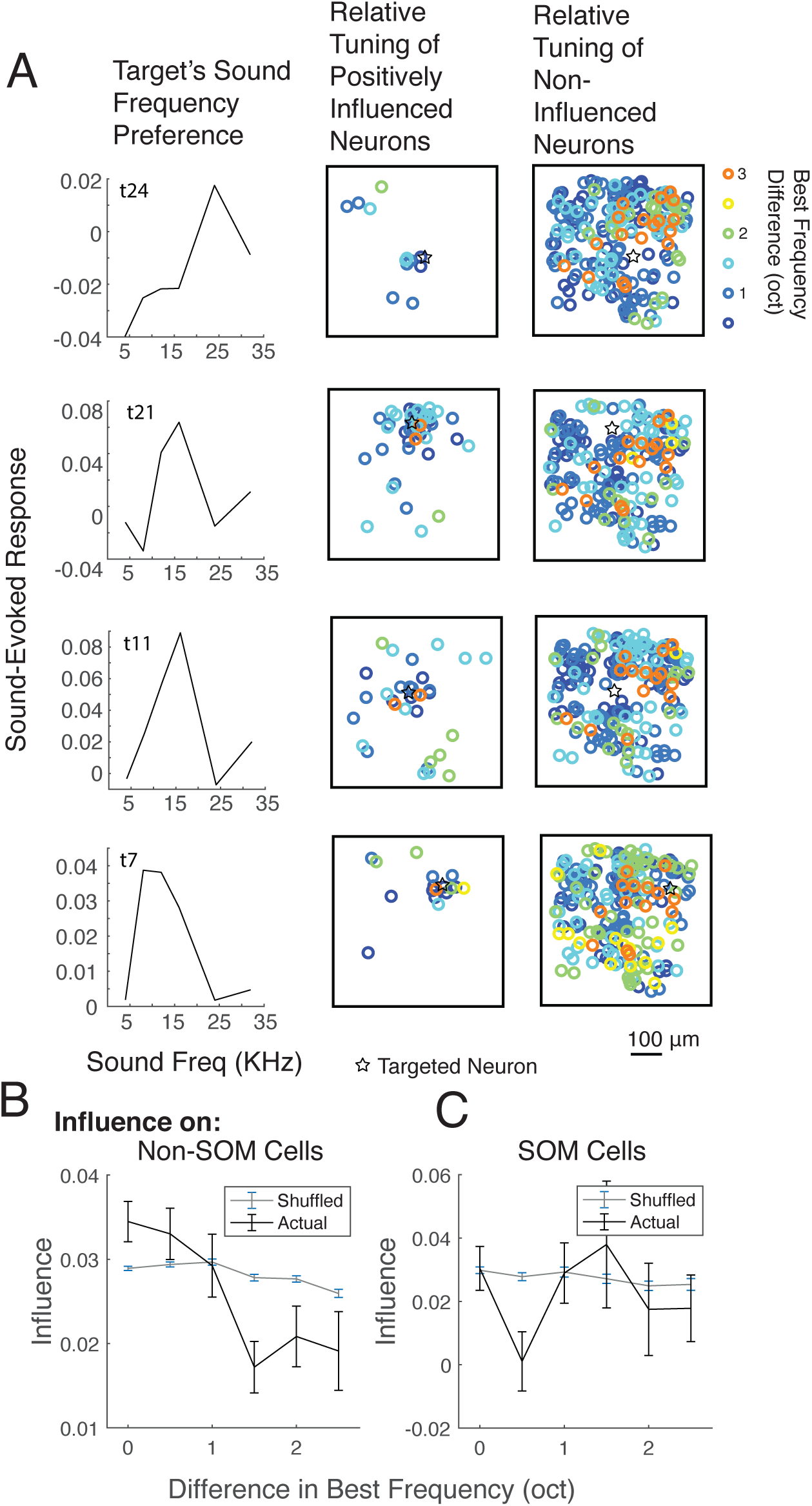
The relationship between E influence and relative sound frequency preference. **(A)** (left) For each sound-responsive and frequency-selective target, we measured its sound frequency preference. We then measured the difference in preferred frequency between the target and all positively influenced (middle) and non-influenced (right) neurons. Each row shows an example sound responsive target and relative tuning of its influence and non-influence maps, all from the same imaging session. Target locations are indicated with black stars in the influence maps. **(B)** Mean influence binned by difference in best frequency between the target and Non-SOM neurons (black). For comparison to an unorganized influence map, the sound frequency preferences were randomly shuffled among the neurons before comparing influence to relative best frequency (grey). **(C)** As in B, for SOM neurons.

## Discussion

In the current study, we set out to compare the spatial structure of local excitatory influence in sensory and association cortex. We targeted individual excitatory (E) neurons with photostimulation while simultaneously monitoring activity in the neighboring population of E and somatostatin-expressing inhibitory interneurons (SOM). While our method cannot reveal the synaptic path from the targeted neuron to the neurons that are modulated, it does allow us to depict the network-level impact of the targeted neuron’s activation. The spatial profile of the local population’s response was similar in AC and PPC, with a highly concentrated positive influence centered on the site of stimulation that decayed quickly over distance, and a more spatially diffuse negative influence that spread farther. However, across areas, the positive influence of E neurons spread farther in SOM neurons in PPC than in AC.

The center-surround organization of local influence that we observed is consistent with known organization of excitatory and inhibitory microcircuits in cortex. The probability of monosynaptic connection between two excitatory neurons decays rapidly with distance (13, 39–42). Single neuron stimulation *in vivo*, where a network-level measurement of a neuron’s influence can be made, similarly reveals that the influence of single excitatory neurons decays with anatomical distance across the local population (7, 8). The inhibitory ‘surround’ triggered by single E cells has a similar spatial pattern across studies, and SOM interneurons may be a key contributor to this surround organization, as SOM neurons receive lateral excitatory synapses from more spatially distant neurons (1, 21). However, it was previously unknown whether the spatial scale of these local circuits is conserved across the cortical hierarchy.

In vivo recordings of large populations of neurons have demonstrated that, like the probability monosynaptic connections, pairwise noise correlations also decay with anatomical distance. However, this decay is more gradual in association-level cortices, especially in SOM neurons (27, 43), suggesting the possibility that the spatial scale of intrinsic connectivity may also be wider in association cortex. Our current study’s results provide some support for this, as the influence on local SOM neurons decayed more slowly over distance, especially in PPC.

Anatomical distance is not the sole determinant of connection probability between two neurons in the cortex. Neurons with similar tuning in V1 have a higher probability of making stronger, bidirectional synaptic connections with each other (5, 6, 14). Here, our results support the possibility that local connectivity is also functionally specific among E cells in auditory cortex, as single E neurons positively influenced the activity of neighboring E neurons with similar sound frequency tuning. Influence on SOM neurons, however, was random with respect to their frequency tuning.

Single neuron resolution optogenetics in visual cortex have also revealed that excitatory neurons tend to influence the local population with functional specificity, though under different stimulation parameters and network conditions these interactions can be either suppressive (8) or enhancing (7). While we did observe a similar spatial pattern of suppressive impacts as these studies, we did not observe suppression that was as dominant, nor did we observe suppressive effects on neurons with similar tuning. Under our study’s experimental conditions (no sensory stimuli, ‘spontaneous’ behavior), suppressive effects may be less prevalent. It will be important in future work to relate the sign of influence to the network state, by presenting stimuli of differing strength and in different behavioral states, such as during task performance.

## Methods

### Experimental Model and Subject Details

All procedures were approved by the University of Pittsburgh Institutional Animal Care and Use Committee. Homozygous SOM-Cre mice (Sst-IRES-Cre, JAX Stock #013044) were crossed with homozygous Ai14 mice (RCL-tdT-D, JAX Stock #007914) obtained from Jackson Laboratory, ME, USA, and all experiments were performed in the F1 generation, which expressed tdTomato in SOM+ neurons. Mice were group housed in cages with between 2 and 4 mice. Adult (8-24 weeks) male and female mice were used for experiments (4 male, 2 female). Mice were housed on reversed 12 hr light/dark cycle, and all experiments were performed in the dark (active) phase.

### Method Details

#### Surgery

Mice were anesthetized with isoflurane (4% for induction, and 1-2% maintenance during surgery), and mounted on a stereotaxic frame (David Kopf Instruments, CA). Ophthalmic ointment was applied to cover the eyes (Henry Schein, NY). Dexamethasone was injected 12-24 hours prior to surgery, and carprofen and dexamethasone (Covetrus, ME) were injected subcutaneously immediately prior to surgery for pain management and to reduce the inflammatory response. Two 2 x 2 mm craniotomies were made over left AC (centered on the temporal ridge, with the posterior edge along the lambdoid suture) and PPC (centered at 2 mm posterior and 1.75 mm lateral to bregma). 1-4 evenly spaced ∼60 nl injections of a viral mixture, containing AAV1-synapsin-l-GCamp6f (Addgene, MA stock #100837 (44)) and AAV9-CAMKII-mScarlet-C1V1-KV2.1 (Addgene, MA stock # 124650) mixed 1:1 and diluted to a titer of ∼1x10^12^ vg/mL using sterile PBS, and were made in each cranial window, centered on the same coordinates listed above. A micromanipulator (QUAD, Sutter, CA) was used to target injections ∼250 μm under the dura at each site, where ∼60 nl virus was pressure-injected over 5-10 minutes. pAAV-CamKIIa-C1V1(t/t)-mScarlet-KV2.1 was a gift from Christopher Harvey (Addgene viral prep # 124650-AAV9; http://n2t.net/addgene:124650 ; RRID:Addgene_124650) (8). Pipettes were not removed until 5 minutes post-injection to prevent backflow. Dental cement (Parkell, NY) sealed a glass coverslip (3mm) over a drop of Kwik Sil (World Precision Instruments, FL) over the craniotomy. Using dental cement, a one-sided titanium headplate was attached to the right hemisphere of the skull. After mice had recovered from the anesthesia, they were returned to their home cages, and received oral carprofen tablets (Fisher Scientific, MA) for 3 days post-surgery.

#### Two-photon microscope

Imaging and photostimulation were performed using a resonant scanning two-photon microscope (Ultima 2Pplus, Bruker, WI). Images were collected at a 30 Hz frame rate and 512 x 512 pixel resolution through a 16x water immersion lens (Nikon CF175, 16X/0.8 NA, NY). On separate days, either AC or PPC was imaged at a depth between 150 and 300 μm, corresponding to layers 2/3 of cortex. For AC imaging, the objective was rotated 35-45 degrees from vertical, and for PPC imaging, it was rotated to 5-15 degrees from vertical, matching the angle of the cranial window implant.

Excitation light was provided by a femtosecond infrared laser (Insight X3, Spectra-Physics, CA) tuned to 920 nm. Photostimulation was achieved by controlling a 1045 nm beam (secondary output from the Insight X3) with an independent set of galvanometers. The 920 and 1045 nm beams were combined with a dichroic (Chroma, ZT1040crb, VT). Green and red wavelength emission light was separated through a 565 nm lowpass filter before passing through bandpass filters (Chroma, ET525/70 and ET595/50, VT). PrairieView software (v5.5, Bruker, WI) was used to control the microscope.

#### Photostimulation

Excitatory neurons expressing mScarlet could be distinguished from the tdTomato+ SOM neurons by the localization of the fluorophores as well as by collecting images at 800 nm excitation, at which only mScarlet is excited. In a typical experiment, 20-30 mScarlet+ neurons and were selected as targets, and 5-10 ‘control’ targets not expressing mScarlet were chosen. the secondary pair of galvanometers was used to direct spiral scans of the 1045 nm beam over targeted neurons.

13µm diameter spirals were directed to each target at 250 Hz for 100 ms, at 20-50 mW. Targets were stimulated in pseudorandom order with a 1 s inter-stimulus interval. At least 100 trial repeats were performed, where different pseudorandom orderings were used across trial repeats.

#### Behavioral monitoring

Running velocity was monitored on pitch and roll axes using two optical sensors (ADNS-98000, Tindie, CA) held adjacent to the spherical treadmill. A microcontroller (Teensy, 3.1, Adafruit, NY) communicated with the sensors, demixing their inputs to produce one output channel per rotational axis using custom code. Outputs controlling the galvanometers were synchronized with running velocity using a digital oscilloscope (Wavesurfer, Janelia, VA).

#### Sound stimuli

Either immediately before or following influence mapping, the same field of view was imaged while presenting pure tone stimuli. One magnetic speaker was pointed to the ear contralateral to the imaging hemisphere (MF1-S, Tucker-Davis, FL). Tones were played at 4, 8, 12, 16, 24, and 32 kHz for 1 s each. Sound responsiveness of each neuron was calculated based on the mean z-scored deconvolved activity of each neuron aligned on sound onset.

#### Image Processing

For each field of view, the raw calcium movies concatenated for motion correction, cell body identification, and fluorescence and neuropil extraction. These processing steps were performed using Suite2p 0.9.3 in Python (45). Suite2p first registered images to account for brain motion, and clustered neighboring pixels with similar time courses into regions of interest (ROIs). ROIs were manually curated using the Suite2p GUI, to ensure that only cell bodies as opposed to dendritic processes were included in analysis, based on morphology. Cells expressing tdTomato (SOM cells), were identified using a threshold applied in the Suite2p GUI based on mean fluorescence in the red channel after bleed-through correction applied by Suite2p’s cell detection algorithm, along with manual correction. For each ROI, Suite2p returned a raw fluorescence timeseries, as well as an estimate of neuropil fluorescence that could contaminate the signal. For each cell, we scaled the neuropil fluorescence by a factor by 0.7 and subtracted this timeseries from the ROI’s raw fluorescence timeseries to obtain a neuropil-corrected fluorescence signal for each selected cell.

#### ΔF/F and deconvolution

Once the neuropil corrected fluorescence was obtained for each neuron, we calculated ΔF/F for each cell in each frame by calculating (F-Fbaseline)/Fbaseline for each frame, where F is the fluorescence of a given cell at that frame and Fbaseline was the eighth percentile of that cell spanning 450 frames before and after (∼15s each way, 30 s total). ΔF/F timeseries were then deconvolved to estimate the relative spike rate in each imaging frame using the OASIS toolbox (46). We used the AR1 FOOPSI algorithm and allowed the toolbox to optimize the convolution kernel, baseline fluorescence, and noise distribution. A threshold of .05 a.u. was applied to remove all events with low magnitude from deconvolved activity timeseries. All analyses were performed with both ΔF/F and deconvolved activity and showed the same trends.

#### Pairwise Noise Correlations

Before computing pairwise noise correlations, each neuron’s trial-averaged response to a given target was subtracted from single trial responses in the 333 ms after the target stimulation onset. These values were then correlated between each target neuron and other neuron in the population, but only during trials that the particular target neuron was not stimulated. The goal was to obtain the pairwise correlations of the neurons outside of times that either neuron was stimulated.

#### Influence calculation

Influence of each target on each other neuron was calculated using deconvolved spike rates as well as dF/F, with similar results. The mean pre-stimulus response of each neuron (binned across the 333 ms prior to the target’s stimulation) was subtracted from its post-stimulus response (in the 333 ms immediately following the target’s stimulation onset). This value was then normalized by the standard deviation of this difference across all trials.

#### Histology

After all imaging sessions had been acquired, each mouse was transcardially perfused with saline and then 4% paraformaldehyde. The brain was extracted, cryoprotected, embedded, frozen, and sliced. Once slide mounted, we stained brains with DAPI to be able to identify structure. We used anatomical structure to verify the locations of our injections in AC and PPC.

#### Quantification and Statistical Analysis

Unless otherwise stated, pairwise comparisons were done with two-sided paired or unpaired permutation (i.e. randomization) tests with 10,000 iterations, where p<.0001 indicates the highest significance achievable given the number of runs performed. All permutation tests were performed for differences in means. When multiple comparisons were made between groups, p values were Bonferroni-corrected.

To compare the effects of brain region, cell type, and inter-somatic distance on influence, we performed a three-way ANOVA. Posthoc tests were calculated as multiple comparisons using the Tukey-Kramer test.

Full values and statistical details from all figures are thoroughly summarized in Tables S1-S3.

## Acknowledgments

We thank members of the Runyan lab for comments on the manuscript. We thank the GENIE project (Janelia) for making GCaMP sensors available for use. We thank the developers of Suite2P and Wavesurfer. This work was supported by the Andrew W. Mellon Predoctoral Fellowship, NIH Predoctoral Training Grant in Basic Neuroscience (T32 NS007433-24, NS007433-21), Pew Biomedical Scholars Program, the Searle Scholars Program, the Klingenstein-Simons Fellowship Award in Neuroscience, and NIH grants NIMH DP2MH122404, NINDS R01NS121913, NIH Supplement NIMH 3DP2MH122404-01S1.

